# Persistent Activation of Chronic Inflammatory Pathways in Long Covid

**DOI:** 10.1101/2024.05.11.593709

**Authors:** Malika Aid, Katherine McMahan, Nicole P Hachmann, Jessica Miller, Erica Borducchi, David Hope, Marjorie Rowe, Eleanor Schonberg, Siline Thai, Ai-ris Y. Collier, Janet Mullington, Dan H. Barouch

## Abstract

Long Covid, or Post-Acute Sequelae of COVID-19 (PASC), involves a spectrum of chronic symptoms following resolution of acute SARS-CoV-2 infection. Current hypotheses for the pathogenesis of Long Covid include persistent SARS-CoV-2, activation of other viruses, tissue damage, autoimmunity, endocrine insufficiency, immune dysfunction, and complement activation. We evaluated 142 participants, including uninfected controls (N=35), acutely infected individuals (N=54), convalescent controls (N=25), and Long Covid patients (N=28), by comprehensive immunologic, virologic, transcriptomic, and proteomic analyses. Long Covid was characterized by persistent inflammatory pathways compared with convalescent controls and uninfected controls, including upregulation of IL-6 and JAK-STAT pathways as well as activation of coagulation, complement, metabolism, and T cell exhaustion pathways. Moreover, robust activation of these pathways during acute COVID-19 infection correlated with the subsequent development of Long Covid. In an independent validation cohort (N=47), Long Covid patients had higher levels of plasma IL-6R compared with convalescent controls and uninfected controls. These data demonstrate that Long Covid is characterized by persistent activation of chronic inflammatory pathways, suggesting novel therapeutic targets and biomarkers of disease.

## Introduction

Long Covid, also known as post-acute sequelae of SARS-CoV-2 infection (PASC), is characterized by a variety of symptoms that can persist for months after recovery from acute COVID-19 infection (1–4). Studies suggest that 10% to 30% of individuals infected with SARS-CoV-2 may experience some form of Long Covid (1, 2, 4, 5). Risk factors for Long Covid include severity of the acute infection, age, gender, and pre-existing health conditions. The most common symptoms are fatigue, shortness of breath, cognitive dysfunction, and joint pain (1, 5). There are currently no effective specific therapies for Long Covid.

The pathophysiology of Long Covid remains unclear but may involve increased complement activation, coagulation, metabolomic abnormalities, endocrine insufficiency, inflammatory responses, and uncoordinated immune responses (1, 6–15). Current hypotheses include persistent virus or viral remnants, autoimmunity, cortisol insufficiency, latent herpesvirus reactivation, T cell dysregulation, and inflammatory tissue damage (3, 5, 7, 9–12, 16–20). Given the diversity of signs and symptoms, treatment is symptomatic and personalized, with an emphasis on rehabilitation.

In this study, we evaluate the immunologic and inflammatory responses in people with Long Covid compared with convalescent controls at 3-6 months and >6 months after initial COVID-19 infection, using immunologic assays, virologic assays, transcriptomics, and proteomics. Acutely infected individuals and uninfected participants served as additional controls. Our data demonstrate that persistent chronic inflammation and immune dysregulation are key features of Long Covid, suggesting novel therapeutic strategies and targets.

## Results

### Study participants

We evaluated samples from 142 participants who were enrolled in the observational biorepository cohorts of the Massachusetts Consortium for Pathogen Readiness and Beth Israel Deaconess Medical Center in Boston, MA. These samples included uninfected controls (Uninfected; N=35), acutely infected individuals <1 month following COVID-19 infection (Acute; N=54), convalescent controls (CC; N=24), and Long Covid patients (LC; N=28) (**Table 1**). Peripheral blood mononuclear cells (PBMC) were collected at 3-6 months (LC: N=25; CC: N=25) and >6 months (LC: N=19) following COVID-19 infection. Plasma samples were similarly collected at 3-6 months (LC: N=23; CC: N=8) and >6 months (LC: N=16; CC: N=6) following COVID-19 infection. A subset of the LC and CC participants also had PBMC (LC: N=4; CC: N=2) and plasma (LC: N=11; CC: N=4) samples from acute COVID-19 infection.

**Table 1.**
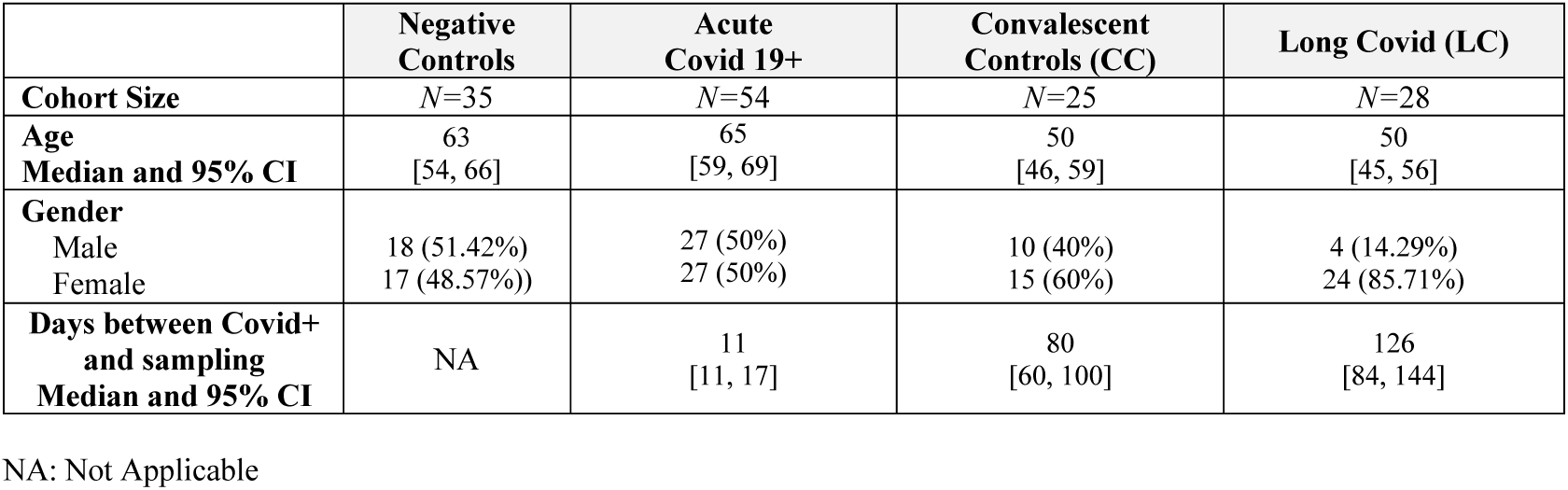
Demographics.

We performed immunologic assays, virologic assays, bulk RNA sequencing, and plasma proteomics to generate comprehensive immune profiles in these individuals (**Fig. 1a**). Clinical symptoms in the Long Covid (LC) cohort included persistent fatigue, shortness of breath, exercise intolerance, brain fog, and abnormal smell and taste **(Fig. 1b)**, similar to prior reports (3, 5, 21–24).

**Fig. 1.**
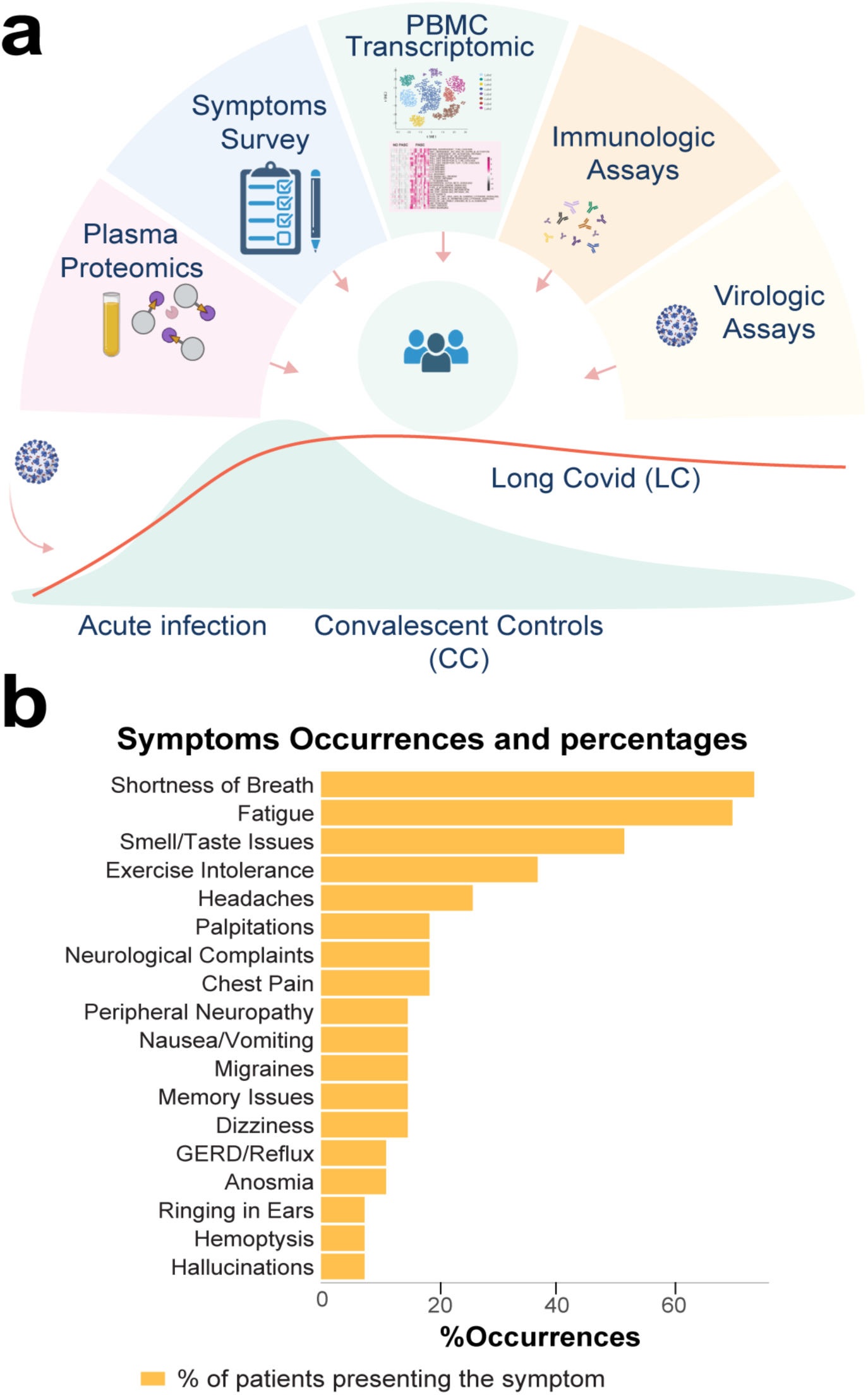
Study overview. **a**, Schematic of samples collection and assays. **b,** Bar plots indicating symptoms occurrences in the long covid group.

### Immunologic and virologic assays

We first assessed SARS-CoV-2 neutralizing antibody (NAb) responses using pseudovirus neutralization assays and T cell responses by pooled peptide interferon-γ ELISPOT assays against SARS-CoV-2 WA1/2020, Delta, and Omicron BA.1 in the CC and LC groups. We did not detect differences in NAb titers, but we observed higher Spike-specific ELISPOT responses to all three variants in the LC individuals compared with the CC individuals (P=0.015, P=0.006, P=0.002, respectively) (**Fig. 2a-b, Supplementary** Fig. 1). The higher cellular immune responses in LC compared with CC individuals may reflect either more severe acute infection or could represent potentially persistent virus (19, 25, 26). However, we did not detect plasma SARS-CoV-2 in any CC or LC individuals by RT-PCR genomic or subgenomic viral load assays (**Fig. 2c)**.

**Fig. 2.**
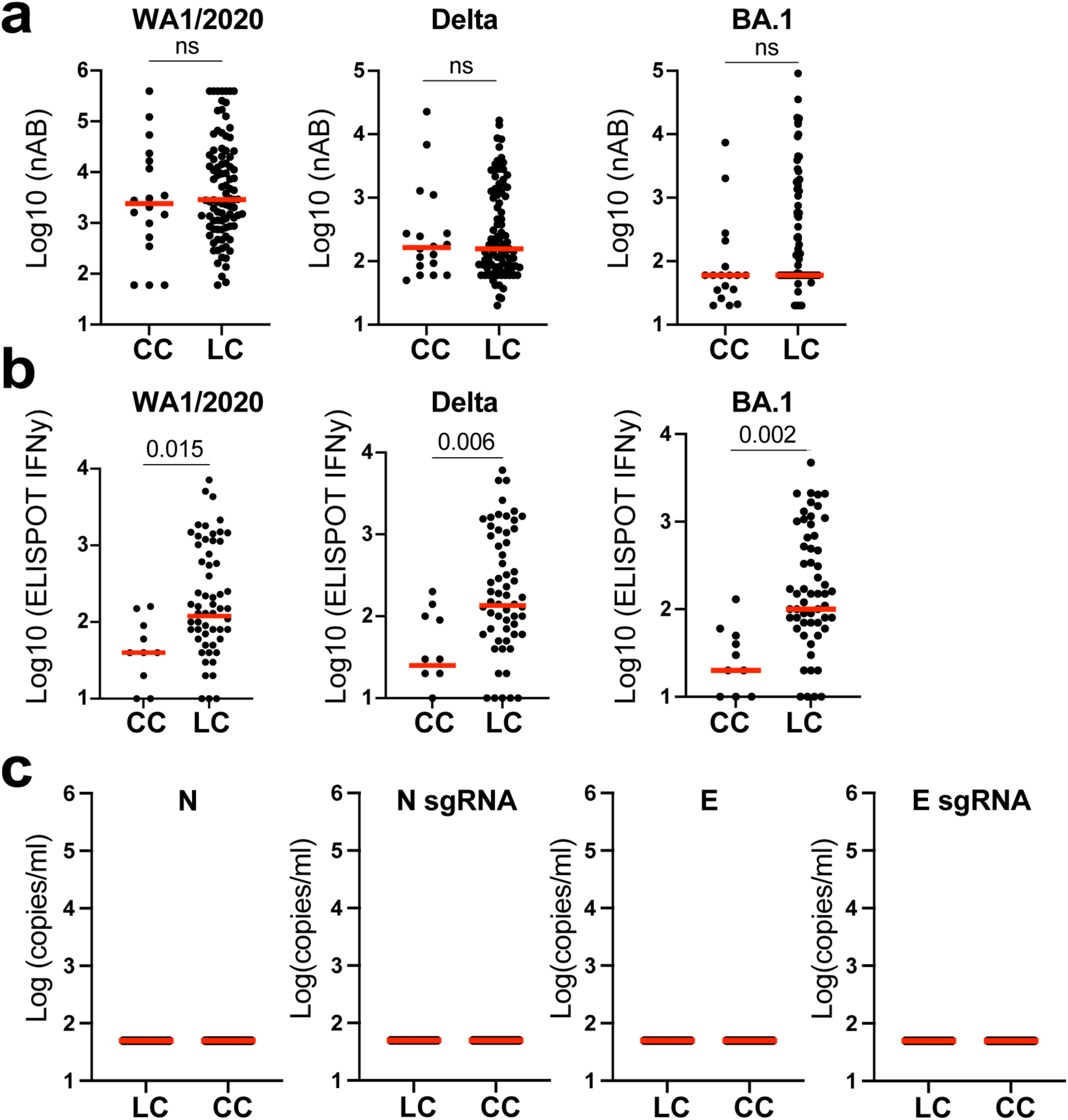
Neutralizing antibody titers, Spike-specific IFN-γ responses, and viral loads in the CC and the LC groups. Neutralizing antibody (NAb) titers (**a**), peptide Spike-specific IFN-γ responses (**b**), and SARS-CoV-2 N and E genomic and subgenomic RNA levels (**c**) in Long Covid (LC) patients and convalescent controls (CC) after Covid-19 infection. Responses were measured against SARS-CoV-2 WA1/2020, Delta, and Omicron BA.1. Medians (red bars) are shown. Groups comparison was performed using the two-sided Mann-Whitney tests, and P values are shown. ns refers to P > 0.05.

### Transcriptomics profiling

We next performed bulk RNA sequencing in PBMC from LC (n=28), CC (n=25), uninfected controls (n=35), and acute COVID-19 (n=54) individuals. Sequenced reads were mapped to the human genome as well as to 20 different viruses. We did not detect differences in viral reads for multiple viruses in the LC compared with CC groups, including human cytomegalovirus (HCMV) and Epstein-Barr virus (EBV) (**Supplementary** Fig. 2). Unsupervised cluster analysis using the whole transcriptome revealed a separation between the LC and other groups, while the CC group clustered with the uninfected controls (**Fig. 3a**). Differential gene expression analysis across these groups showed upregulation of multiple inflammatory markers in the LC compared with the CC groups and uninfected controls, including the signaling molecules LIFR and JAK2; chemokines CXCL2, CXCL3, and CCL3; cytokines IL10, NLRP3, IFNG, IL6, TNF, IL1B, IL1A; and complement and coagulation proteins C5, F3, and THBS1 (**Fig. 3b-c, Supplementary** Fig. 3a-b). Genes associated with natural killer function such as KLRC4, KLRC1, KLRC2 and transcription factors such LEF1, GATA6, BACH2 were decreased in the LC group compared with the CC and uninfected control groups (**Fig. 3d, Supplementary** Fig. 3a-b).

**Fig. 3.**
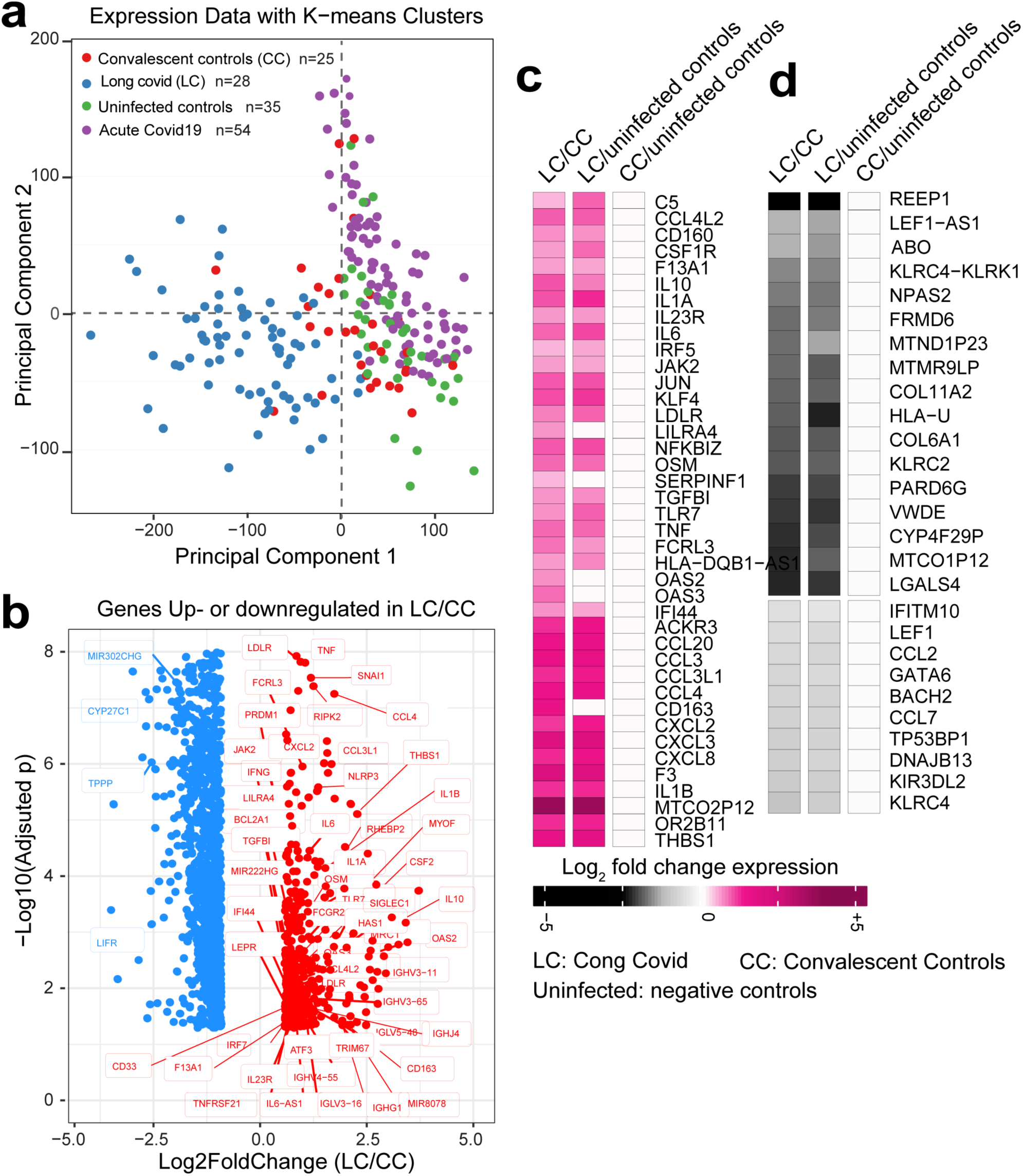
PBMC bulk RNA sequencing in long covid (LC), convalescent controls (CC), acute covid-19+ and uninfected controls. **a**, K-mean clustering using the whole transcriptome and all groups. **b**, Scatter plot of the top genes upregulated (in red) or downregulated (in blue) in LC compared with CC. X-axis: log2 fold change expression (LC/CC). Y-axis: –Log10 (adjusted P, BH method). **c-d,** Top proinflammatory genes decreased or increased in LC compared with CC. Heatmap shows the normalized mRNA expression ranging from blue (down regulated) to red (upregulated).

We performed Gene Set Enrichment Analysis (GSEA, FDR q-value<0.05) (27) to compare the expression profiles in PBMC of the LC and CC groups at 3-6 months following acute COVID-19 infection. The LC group was characterized by higher levels of innate immune cell signatures for monocytes, macrophages, neutrophils and dendritic cells; complement and coagulation cascade signatures; and cytokine and chemokine signaling pathways, including IL6, IL8, IL10, IL12, IL17, JAK_STAT, and type I interferon pathways (**Fig. 4a**) (6, 7, 20, 21, 28, 29). We also observed increased metabolic activity in the LC group compared with the CC group involving amino acid and lipid metabolism (**Fig. 4a**), consistent with prior reports (6, 14, 15, 20, 29). A smaller number of pathways were downregulated in the LC group compared with the CC group, including RNA processing, ribosome, and spliceosome signaling (**Fig. 4b**). The pathways that showed the most significant enhancement in the LC group compared with the CC group were the IL6, JAK_STAT, IL1R, mast cell, coagulation, complement, bile acid metabolism, ascorbate/aldarate, and leptin signaling (**Fig. 4c**). No differences in these pathways were observed between the CC and uninfected controls (**Fig. 4c**).

**Fig. 4.**
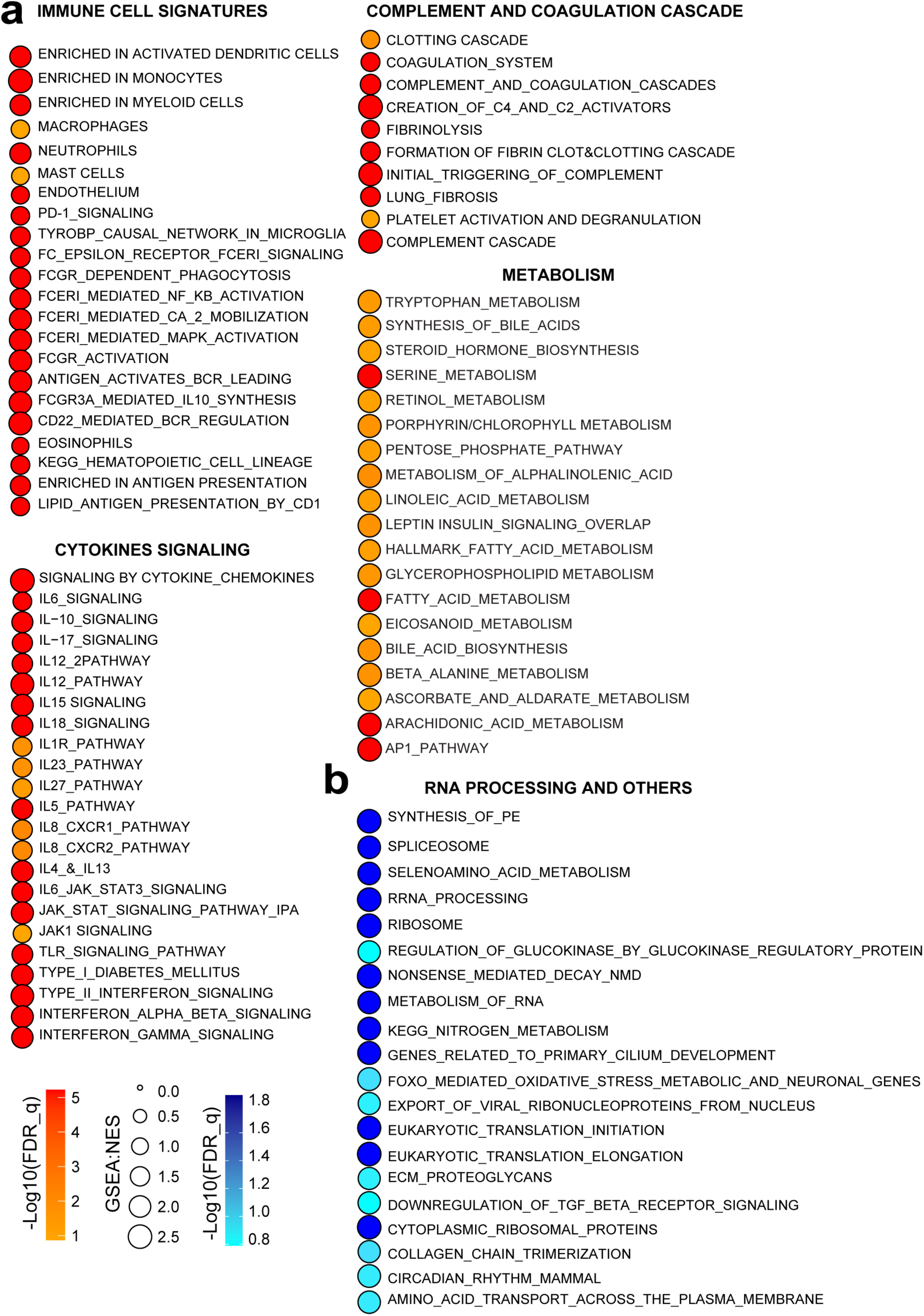
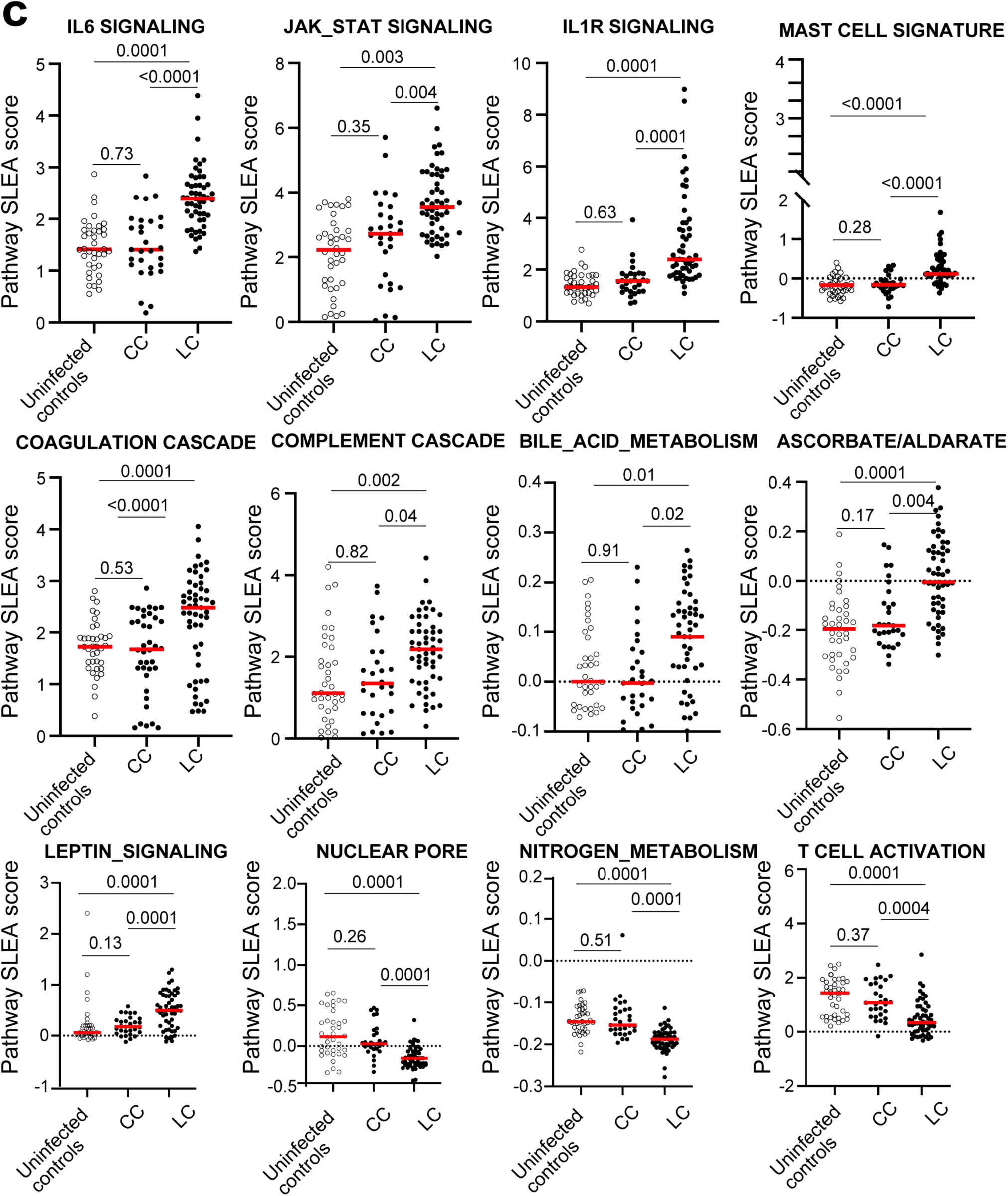
Pathways upregulated (red) or downregulated (blue) in the LC (*n=28*) compared with CC (*n=25*) groups at 3-6 months after COVID-19 infection in PBMC. **a-b**, Circle plots of pathways normalized enrichment scores measured by gene set enrichment analysis (GSEA). Increased pathways shown in red (a) and decreased pathways shown in blue (b) gradients. Circle size: pathways normalized enrichment score. **c,** Single sample GSEA (ssgsea) scores of pathways increased or decreased in LC compared with CC and uninfected controls. Each dot represents a sampling time point. Groups comparison was performed using the two-sided Mann-Whitney tests, and P values are shown. Medians (red bars) are shown.

We observed downregulation of certain T cell pathways in the LC group compared with the CC group (**Fig. 4c**), including T cell differentiation and activation pathways and T regulatory cell signatures (**Fig. 5a-b**). However, signatures of T cell exhaustion and PD-1 signaling pathways were significantly increased in the LC group compared with the CC group (**Fig. 5b**), suggesting that Long Covid is associated with T cell dysfunction. T cell signaling correlated inversely with IL6, JAK-STAT, and interferon signaling (**Fig. 5c**).

**Fig. 5.**
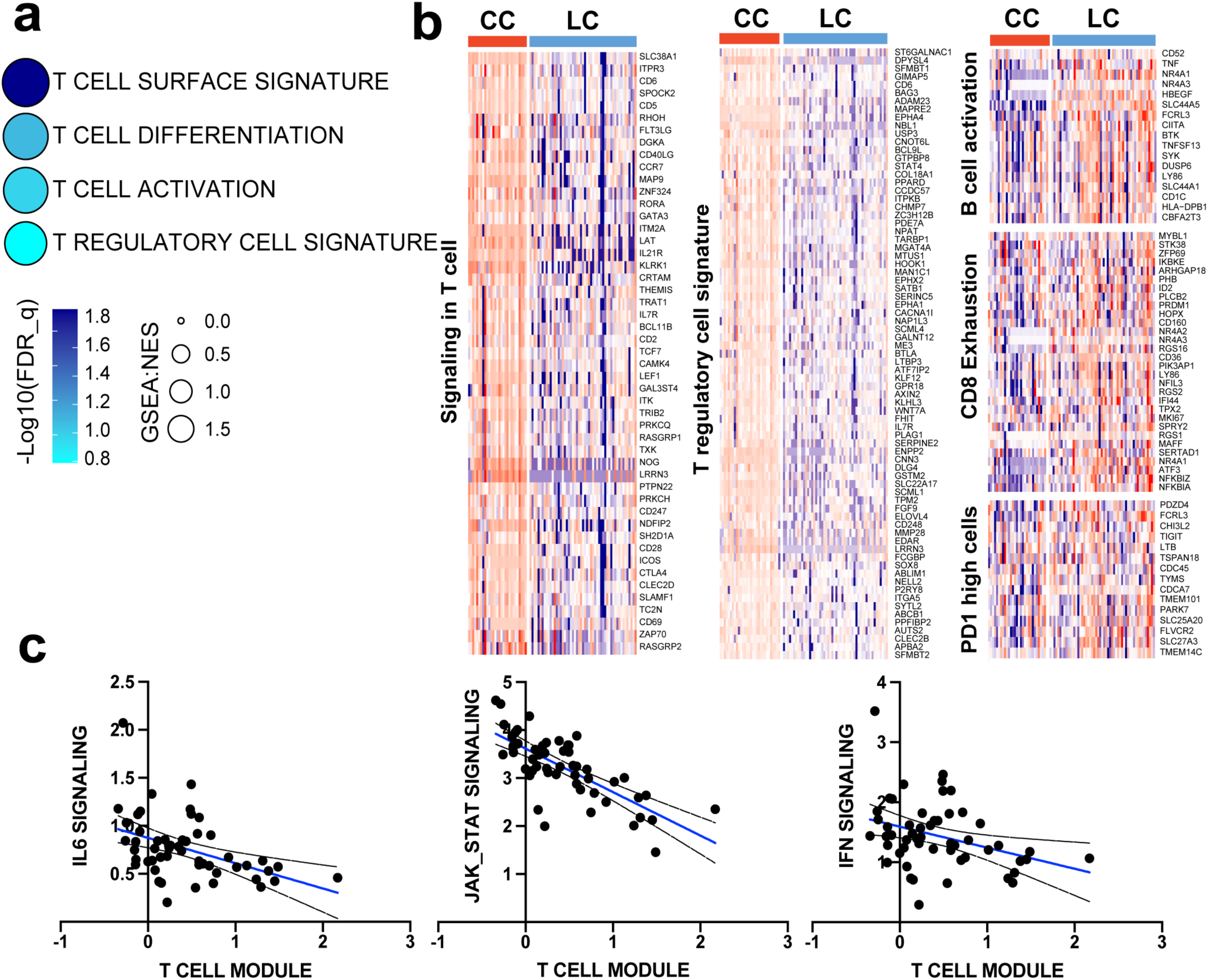
Signatures of T cell signaling in the LC and CC groups in PBMC by bulk RNA sequencing 3-6 months after COVID-19 infection. **a**, Circle plots of pathways normalized enrichment scores measured by gene set enrichment analysis (GSEA). Increased pathways shown in red and decreased pathways shown in blue gradients. Circle size: pathways normalized enrichment score. **b,** T regulatory and T cell signaling; B cell, Exhausted CD8 and PD1 signatures increased or decreased in LC compared with CC. **c**, Regression analyses of ssGSEA pathway score of T cell signatures correlated with IL6, JAK_STAT and interferon signaling pathways. Each point represents an individual sampling time point (3-6 months) along with corresponding 95% confidence intervals (CI) which illustrate the precision of these estimations. P-values were calculated using two-tailed t-test, indicating the statistical significance of the observed associations (p<0.001).

Additional signatures associated with autoimmune diseases were also increased in the LC group compared with the CC group (**Supplementary** Fig. 3c). Moreover, proinflammatory signatures that were increased in the LC group were co-correlated with each other (**Supplementary** Fig. 3d) and with the IL6 signaling signature (**Supplementary** Fig. 3e).

### Proteomics profiling

Plasma proteomics revealed similar differences between the LC and CC groups, consistent with the transcriptomic data (**Fig. 6**). We observed increased levels of immune cell signatures, such as neutrophils, monocytes and epithelial cells; cytokine signaling, including IL6, IL12, IL8, NFKPB, JAK-STAT, and corticotropin releasing hormone; complement and coagulation cascades; and certain metabolic pathways in the LC group compared with the CC group (**Fig. 6a, c, Supplementary** Fig. 4). In contrast, pathways of cytotoxic T cell and NK_DCs cross talk signaling were downregulated in the LC group compared with the CC group (**Fig. 6b, c**). IL6 signaling in plasma inversely correlated with cytotoxic T cell, amino acid metabolism, and DNA damage pathways and directly correlated with metabolic pathways (**Supplementary** Fig. 5).

**Fig. 6.**
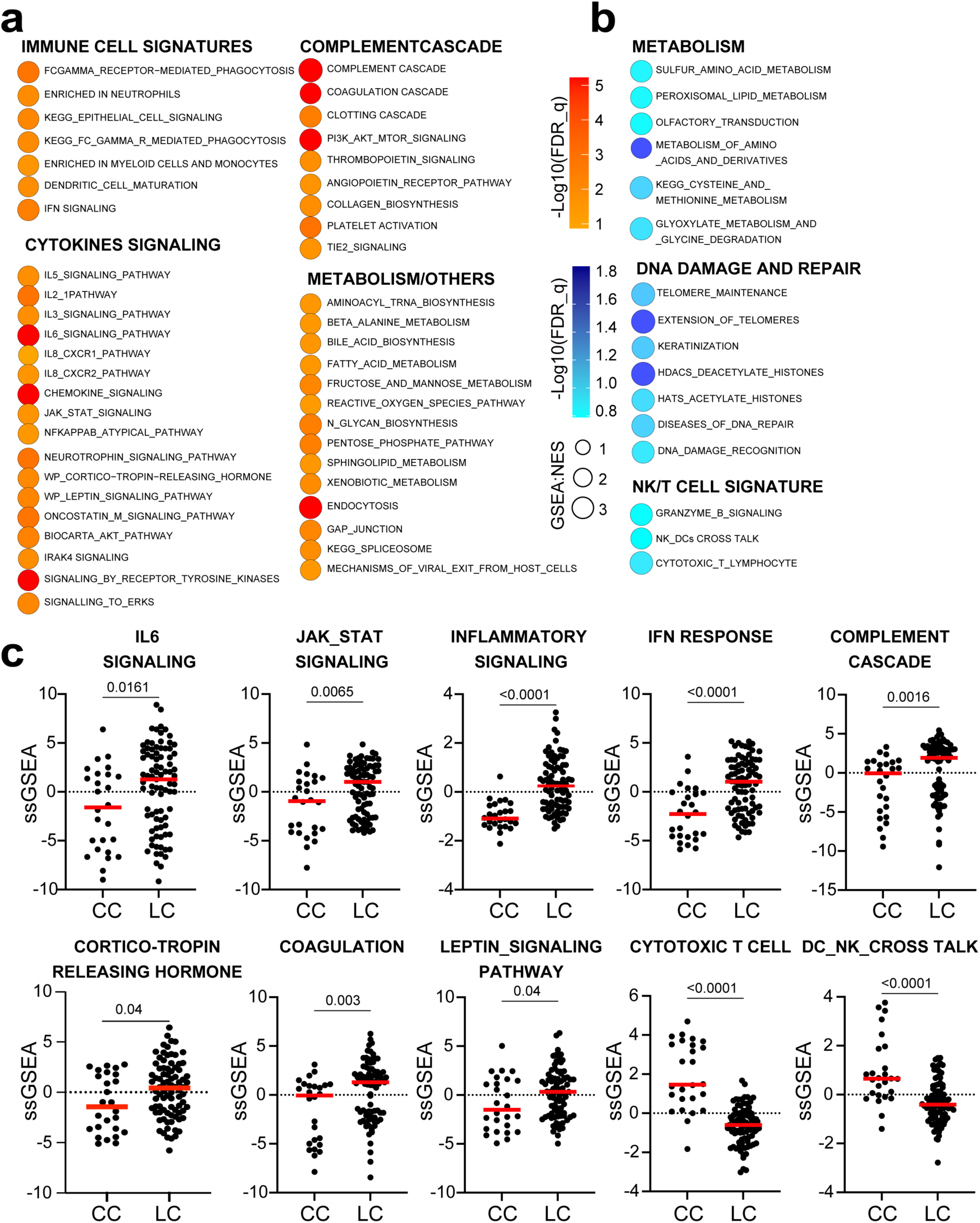
Pathways upregulated (red) or downregulated (blue) in the LC (*n=24*) compared with the CC (*n=9*) groups 3-6 months after COVID-19 infection in plasma by proteomics profiling. **a-b**, Circle plots of pathways normalized enrichment scores measured by gene set enrichment analysis (GSEA, nominal p<0.05) in LC vs. CC. Increased pathways shown in red (**a**) and decreased pathways shown in blue (**b**) gradients. Circle size: pathways normalized enrichment score. **c,** Single sample GSEA (ssgsea) scores of pathways increased or decreased in LC compared with CC and uninfected controls. Each dot represents a sampling time point. Groups comparison was performed using the two-sided Mann-Whitney tests, and P values are shown. Medians (red bars) are shown.

In a pilot study, we assessed whether there were differences in individuals during acute COVID-19 infection who subsequently developed Long Covid or became convalescent controls in the subset of participants in the LC and CC groups who had samples both <1 month and >3 months after COVID-19 infection. During acute COVID-19 infection, increased IL6, complement, and cortisol signaling correlated with the subsequent development of Long Covid at 3-6 and >6 months (**Supplementary Fig. 6a-b, Supplementary Fig. 7**). Feature importance analyses revealed that plasma level of IL6R, IL6ST, STAT1, STAT3, PTPN11 and MAPK1,3,4 during acute COVID-19 infection (< 1month) was associated with the subsequent development of LC at 3-6 and >6 months (**Supplementary Fig. 6c**). These data suggest that upregulation of these pathways during acute COVID-19 infection may contribute to the pathogenesis of Long Covid.

### Validation studies

To validate our findings in an independent cohort, we evaluated plasma levels of proinflammatory cytokines and chemokines and found increased levels of IL6R in individuals with LC (n=19) compared with CC (n=13) and uninfected controls (n=13), using both and ELISA and a meso-scale discovery (MSD) assay at 3-6 months and >6 months after COVID-19 infection (**Fig. 7a-b)**. These findings support the link between chronic inflammation and Long Covid (7, 10, 17, 29) and suggest that plasma inflammatory markers should be explored as potential biomarkers for Long Covid.

**Fig. 7.**
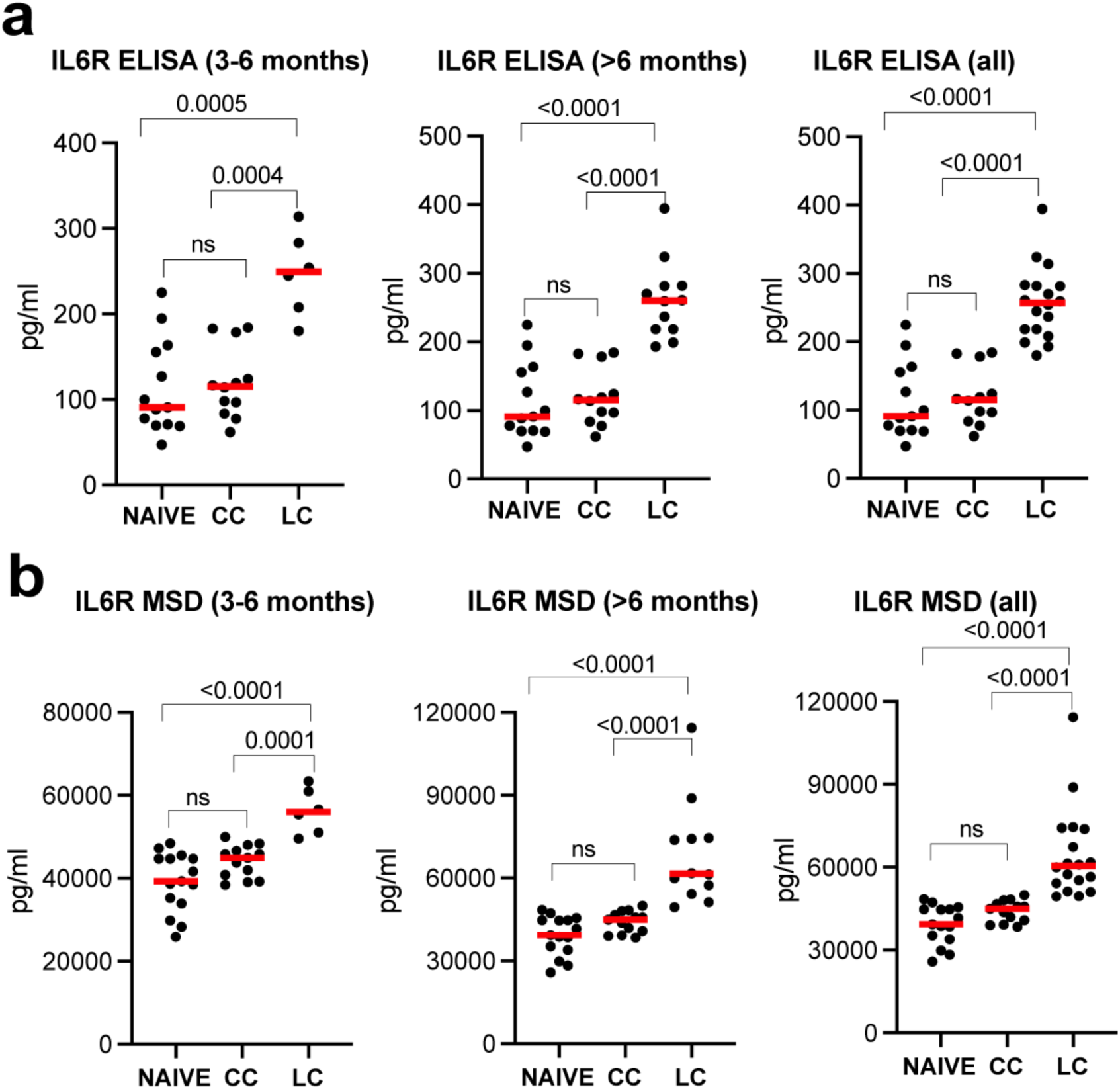
Plasma IL6R upregulation in a validation cohort by ELISA and MSD assays. IL6R plasma level measured by ELISA (**a**) and mesoscale discovery (MSD) assays (**b**) in LC (n=19), CC (n=13), and naïve uninfected controls (n=13). Comparative analysis of IL6R levels measured through MSD and ELISA techniques in subjects across three groups: Uninfected, CC, and LC groups. Mann-Whitney U Test was used to assess the statistical significance of differences between the median IL6R levels among groups. The results, illustrated as P-values located above the inter-group comparison lines, indicate a highly significant difference (P < 0.05) in IL6R levels in both detection methods. Medians (red bars) are shown. ns refers to P > 0.05. data were shown for 3-6 months, > 6 months and all samples combined for each assay.

## Discussion

The pathogenesis of Long Covid has remained unclear, hampering efforts to develop specific therapies. In this study, we demonstrate that individuals with Long Covid have persistent activation of pathways of chronic inflammation, cytokine signaling, coagulation and complement cascades, metabolic dysregulation, and immune cell exhaustion compared with convalescent controls. These findings suggest potential therapeutic targets and biomarkers of disease that should be evaluated in future clinical trials.

Activation of innate immune cell signatures, signaling by proinflammatory cytokines such as IL6, IL4, IL5 and IL12 and JAK-STAT signaling, complement and coagulation, and metabolic pathways were observed at both 3-6 months and >6 months after infection in the LC group compared with the CC group. The LC group exhibited reduced pathways for granzyme B and cytotoxic T cell signaling, indicating T cell dysregulation and exhaustion, which is consistent with prior studies (16, 30). We also observed in a pilot study that early activation of IL-6 and complement pathways during acute COVID-19 infection correlated with the subsequent development of Long Covid, suggesting the mechanistic importance of activation of these pathways in the pathogenesis of Long Covid and the potential for early intervention strategies.

Our study confirmed and extended prior reports of metabolic dysregulation in Long Covid (6, 14, 15, 17, 18, 20, 22, 29). We also observed decreased activity of the telomere maintenance pathways and DNA damage recognition pathways in the LC group compated with the CC group, which may signal impaired tissue regeneration in patients with Long Covid (31, 32).

In conclusion, our data demonstrate that Long Covid is characterized by persistent chronic inflammation and immune exhaustion. Clinical trials of therapeutic drugs for Long Covid currently focus on antiviral agents such as Paxlovid, which target potential residual viral replication. Our study identifies additional therapeutic targets that should be explored, including chronic inflammation pathways such as the IL6 and JAK-STAT pathways as well as pathways associated with T cell dysfunction. Clinical trials of anti-inflammatory drugs targeting these pathways are planned.

## Acknowledgements

We acknowledge support from the Massachusetts Consortium for Pathogen Readiness, Gates Foundation (INV-027406, INV-041469), and the National Institutes of Health (CA260476) (D.H.B.). Next generation sequencing services were provided by the Emory NPRC Genomics Core (NIH P51 OD011132, S10 OD026799). Proteomics data were generated by the Beth Israel Deaconess Medical Center Proteomics Core.

## Author contributions

This study was designed by M.A and D.H.B. Immunologic and virologic assays were performed by K.M., N.P.H., J.M., E.B., D.H. The validation cohort was led by M.R., E.S., S.T., A.Y.C., and J.M. The paper was written by M.A. and D.H.B. with all co-authors.

## Data sharing

All data are available in the manuscript or the supplementary material. Transcriptomics raw data has been deposited in the GEO database under accession number GSE226260 (access token is kxezccgcnhcnvmh).

## Correspondence and requests

Correspondence and requests for materials should be addressed to D.H.B.

## Methods

### Neutralizing Antibody Assay

Neutralizing antibody (NAb) titers against SARS-CoV-2 variants were determined using pseudo-typed viruses expressing a luciferase reporter gene. In brief, a luciferase reporter plasmid pLenti-CMV Puro-Luc (Addgene), packaging construct psPAX2 (AIDS Resource and Reagent Program), and Spike protein expressing pcDNA3.1-SARS-CoV-2 SΔCT were co-transfected into human embryonic kidney (HEK)293 T cells (ATCC CRL_3216) with lipofectamine 2000 (ThermoFisher Scientific). Pseudo-typed viruses of SARS-CoV-2 variants were generated using the Spike protein from WA1/2020 (Wuhan/WIV04/2019, GISAID accession ID: EPI_ISL_402124), B.1.617.2 (Delta, GISAID accession ID: EPI_ISL_2020950), and Omicron BA.1 (GISAID accession ID: EPI_ISL_7358094.2). 48 h post-transfection, the supernatants containing the pseudo-typed viruses were collected and purified by filtration with 0.45-μm filter. To determine NAb titers in human sera, HEK293T-hACE2 cells were seeded in 96-well tissue culture plates at a density of 2 × 104 cells per well overnight. Three-fold serial dilutions of heat-inactivated serum samples were prepared and mixed with 60 μl of pseudovirus, and incubated at 37 °C for 1 h before adding to HEK293T-hACE2 cells. 48 h later, cells were lysed in Steady-Glo Luciferase Assay (Promega) according to the manufacturer’s instructions. SARS-CoV-2 neutralization titers were defined as the sample dilution at which a 50 % reduction (NT50) in relative light units was observed relative to the average of the virus control wells. Titers greater than 1:20 were considered positive.

### IFN-γ Enzyme-Linked Immunospot (ELISPOT) Assay

Cellular immune responses specific to SARS-CoV-2 were assessed by interferon-(IFN-γ) ELISPOT assays using pools of overlapping 15-amino-acid peptides for Wisconsin, delta and omicron variants (21^st^ Century Biochemicals, USA). 96-well multiscreen plates (Millipore, MA, USA) were coated with 1µg/well of mouse anti-human IFN-γ (MabTech., USA) overnight in endotoxin-free Dulbecco’s PBS (DPBS) overnight at 4°C. Plates were washed with DPBS 3 times, and blocked using RPMI 1640 media containing 10% FBS for 2-4 hours at 37°C. Peptides pools were prepared at a concentration of 2 µg/well, and 200,000 cells/well were added. Peptides and cells were incubated for 15-20 h at 37°C. The plates were washed with DPBS-Tween 7 times and then incubated with 1µg/mL per well of biotinylated anti-human IFN-γ (MabTech, USA) for 2 to 4 hours at room temperature, followed by 4 washes with DPBS-Tween and 1.33µg/mL per well of alkaline phosphatase-conjugated anti-biotin (Rockland, USA) for 2 to 3 hours at room temperature. Plates were developed with nitroblue tetrazolium-5-bromo-4-chloro-3-indolyl-phosphate chromogen (Pierce, IL, USA), stopped by washing with tap water, and read using an ELISPOT reader (KS ELISPOT Reader, Carl Zeiss). The numbers of spot-forming cells (SFC) per 106 cells were calculated subtracted over background (PBMCs incubated with media and DMSO without peptide).

### Bulk RNA Sequencing

PBMCs were lysed in 700 μL of Trizol and then extracted using the miRNeasy Mini kit (Qiagen) with on-column DNase digestion. RNA quality was assessed using a TapeStation 4200 (Agilent) and then ten nanograms of total RNA was used as input for cDNA synthesis using the Clontech SMART-Seq v4 Ultra Low Input RNA kit (Takara Bio) according to the manufacturer’s instructions. Amplified cDNA was fragmented and appended with dual-indexed barcodes using the Nextera XT DNA Library Preparation kit (Illumina). Libraries were validated by capillary electrophoresis on a TapeStation 4200 (Agilent), pooled at equimolar concentrations, and sequenced with PE100 reads on an Illumina NovaSeq 6000, yielding ∼30 million reads per sample on average. Alignment was performed using STAR version 2.7.3a(^33^)and transcripts were annotated using a composite genome reference which included GRCh38 Ensembl release 100 and SARS-CoV-2 (GCF_009858895.2, ASM985889v3, MN985325.1). Transcript abundance estimates were calculated internal to the STAR aligner using the algorithm of htseq-count(34). Transcript abundance estimates were calculated internal to the STAR aligner using the algorithm of htseq-count(34). DESeq2 was used for normalization(^35^), producing a normalized read count.

### Plasma Proteomics

55 ml serum or plasma from all participants, five pooled plasma controls, and one buffer control were analyzed using the SomaScan® Assay Kit for human plasma V4.1 (Cat#. 900-00021), measuring the expression of 6596 unique human protein targets using 7596 SOMAmer (slow off-rate modified aptamer) reagents, single-stranded DNA aptamers, according to the manufacturer’s standard protocol (SomaLogic; Boulder, CO). The modified aptamer binding reagents, SomaScan Assay, its performance characteristics, and specificity to human targets have been previously described (36). The assay used standard controls, including 12 hybridization normalization control sequences used to control for variability in the Agilent microarray readout process, as well as five human calibrator control pooled plasma replicates and 3 Quality Control (QC) pooled replicates used to mitigate batch effects and verify the quality of the assay run using standard acceptance criteria. The readout is performed using Agilent microarray hybridization, scan, and feature extraction technology.

Twelve Hybridization Control SOMAmers are added alongside SOMAmers to be measured from the serum samples and controls of each well during the SOMAmer elution step to control for readout variability. The control samples are run repeatedly during assay qualification and robust point estimates are generated and stored as references for each SOMAmer result for the Calibrator and QC samples. The results are used as references for the SomaScan V4.1 Assay. Plate calibration is performed by calculating the ratio of the Calibrator Reference relative fluorescence unit (RFU) value to the plate-specific Calibrator replicate median RFU value for each SOMAmer. The resulting ratio distribution is decomposed into a Plate Scale factor defined by the median of the distribution and a vector of SOMAmer-specific Calibration Scale Factors. Normalization of QC replicates and samples is performed using Adaptive Normalization by Maximum Likelihood (ANML) with point and variance estimates from a normal U.S. population. Post-calibration accuracy is estimated using the ratio of the QC reference RFU value to the plate-specific QC replicate median RFU value for each SOMAmer. The resulting QC ratio distribution provides a robust estimate of accuracy for each SOMAmer on every plate. SomaScan RFU values and clinical information are obfuscated to protect personally identifiable information while preserving biologically relevant biomarkers. These reference data sets were provided by SomaLogic. We used the Linear Models for Microarray Data (Limma) R package (37) to identify differentially expressed proteins. The method involves fitting a linear model to the data and then performing a t-test to identify proteins that are differentially expressed between two or more groups. P values were corrected for multiple testing using the Benjamini-Hochberg method. R packages ggplot2, ComplexHeatmap, were used to generated the figures. Pathway enrichment analysis was performed using Gene Set Enrichment Analysis (GSEA) (27) against the C2 database from the Molecular Signatures Database (MSigDB) (38).

### ELISA

Cytokines were assessed by ELISA. An anti-human cytokine coating antibody was adsorbed onto 96-microwell plates. Microwells were washed with wash buffer, and human serum samples and human cytokine standards were prediluted in assay buffer and added to each plate. Plates were then incubated for 1 hour, prior to addition of anti-human cytokine horseradish peroxidase (HRP). Following an additional 1-hour incubation, plates were again washed with wash buffer. SeraCare KPL TMB SureBlue Start solution was added to each well; plate development was halted by adding SeraCare KPL TMB Stop solution to each well. The absorbance at 450nm was recorded with a VersaMax microplate reader (Molecular Devices). The standard curve was prepared from human cytokine standard dilutions and human cytokine concentration determined. For each sample, the cytokine concentration was calculated using a 4-parameter logistic curve fit; cytokine concentrations of the unknown samples are interpolated from linear portion of the standard curve generated from human cytokine standards of known concentration.

### Mesoscale Discovery Assay

Serum levels of Human IL-6R were tested using R-PLEX Human IL-6R kits from MesoScale Discovery (K1510GR-2, MSD, Rockville, MD) by Metabolism and Mitochondrial Research Core (Beth Israel Deaconess Medical Center, Boston, MA) following manufacture’s instruction. In brief, the plate was coated using provided biotinylated capture antibody with diluent 100. The highest Calibrator standard is 2000 pg/mL. Then a 4-fold serial dilution were done to generate 7 calibrator curve using 4-parameter logistic model. The detection limit (LLOD) is 0.4pg/mL. Samples were thawed on ice and diluted in 1:200 ratio using Diluent 7 provided by the kit. The assay plate was read by MESO QUICKPLEX SQ 120 instrument and data were analyzed by Discovery workbench 4.0 software.

### Statistical Analyses

Data from RNAseq, proteomics, were submitted to an exploratory analysis conducted in R environment or in Graphpad PRISM 9.0 software (GraphPad Software, San Diego,California, USA). Enrichment analysis of the canonical pathways of the significantly differentially expressed genes, or proteins was conducted using GSEA. Further statistical tests were done either in R (R Core Team, 2020) or GraphPad PRISM 9.0 software (GraphPad Software, San Diego, California, USA). Group comparisons was evaluated using GraphPad Prism. The analysis delineates a comparative exploration of the groups’ median values, visually represented through the data distribution and median lines. Statistical significance, denoted by the calculated P-value, is highlighted above the comparison line, with conventional thresholds indicating significance: a P-value less than 0.05 is considered statistically significant. The two-sided Mann-Whitney U Test was used in identifying significant differences between the groups under study, without relying on the assumption of data normality. The R random forest package (randomForest) was used to select for the top plasma features, during the acute phase, that are associated with the subsequent development of Long Covid. Features importance scores were measured using the Gini coefficient, where the higher the value of mean decrease Gini score, the higher the importance of the variable in the model.

## Supplementary Figure Legends

**Supplementary Fig. 1.**
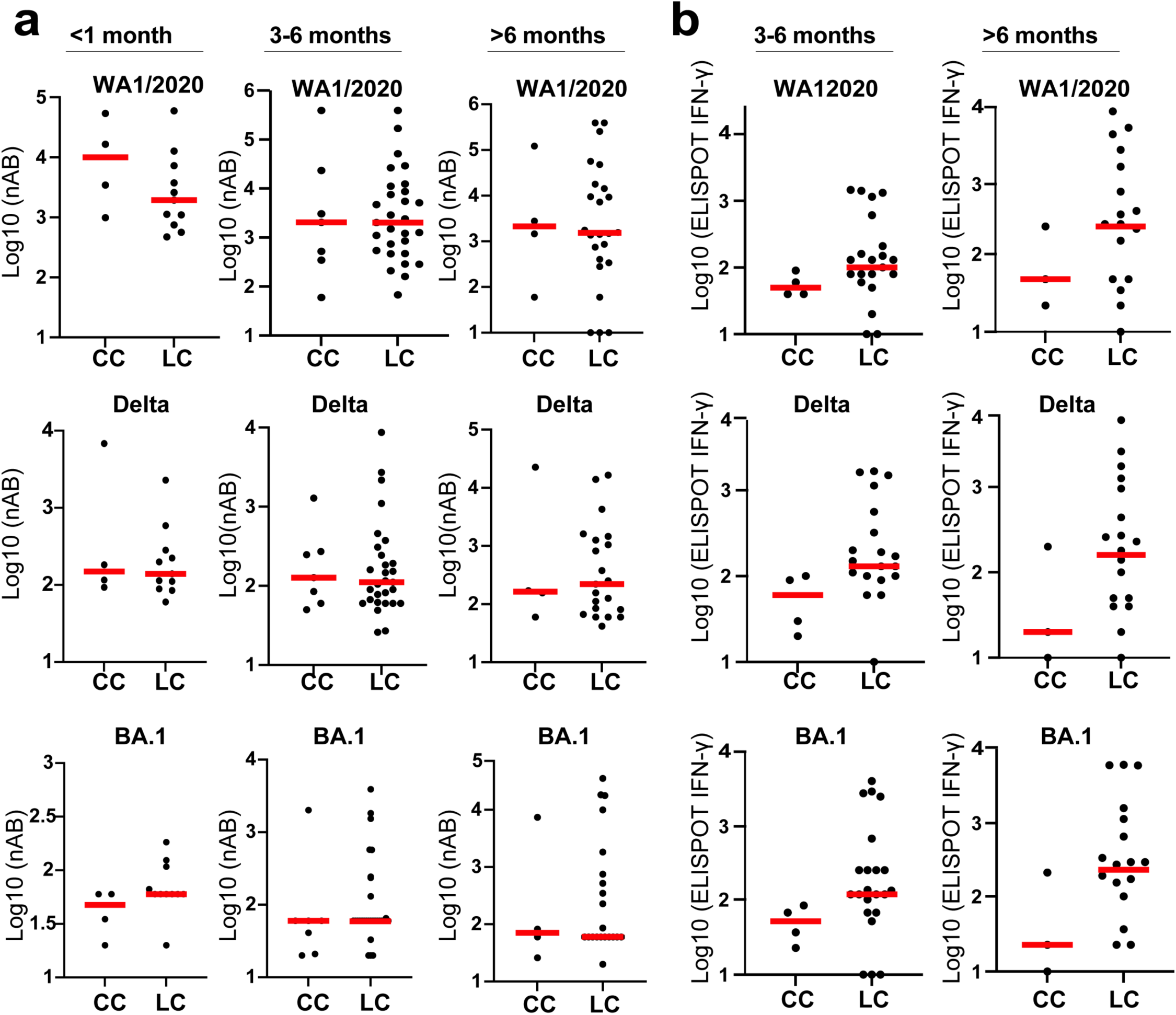
SARS-CoV-2 neutralizing antibody and T cell responses in recovered controls (CC) and long covid (LC) patients. a, Neutralizing antibody (NAb) titers were assessed before 1month, 3-6 months and after 6 months to covid-19 infection in recovered (CC) and long covid patients (LC) by intracellular cytokine staining. Responses were measured against SARS-CoV-2 WA1/2020, Delta and BA.1. Medians (red bars) are shown. **b,** Pooled peptide Spike-specific IFN-g responses were assessed at 3-6 months and after 6 months to covid-19 infection in recovered and long covid patients by intracellular cytokine staining. Responses were measured against SARS-CoV-2 WA1/2020, Delta and BA.1. Medians (red bars) are shown.

**Supplementary Fig. 2.**
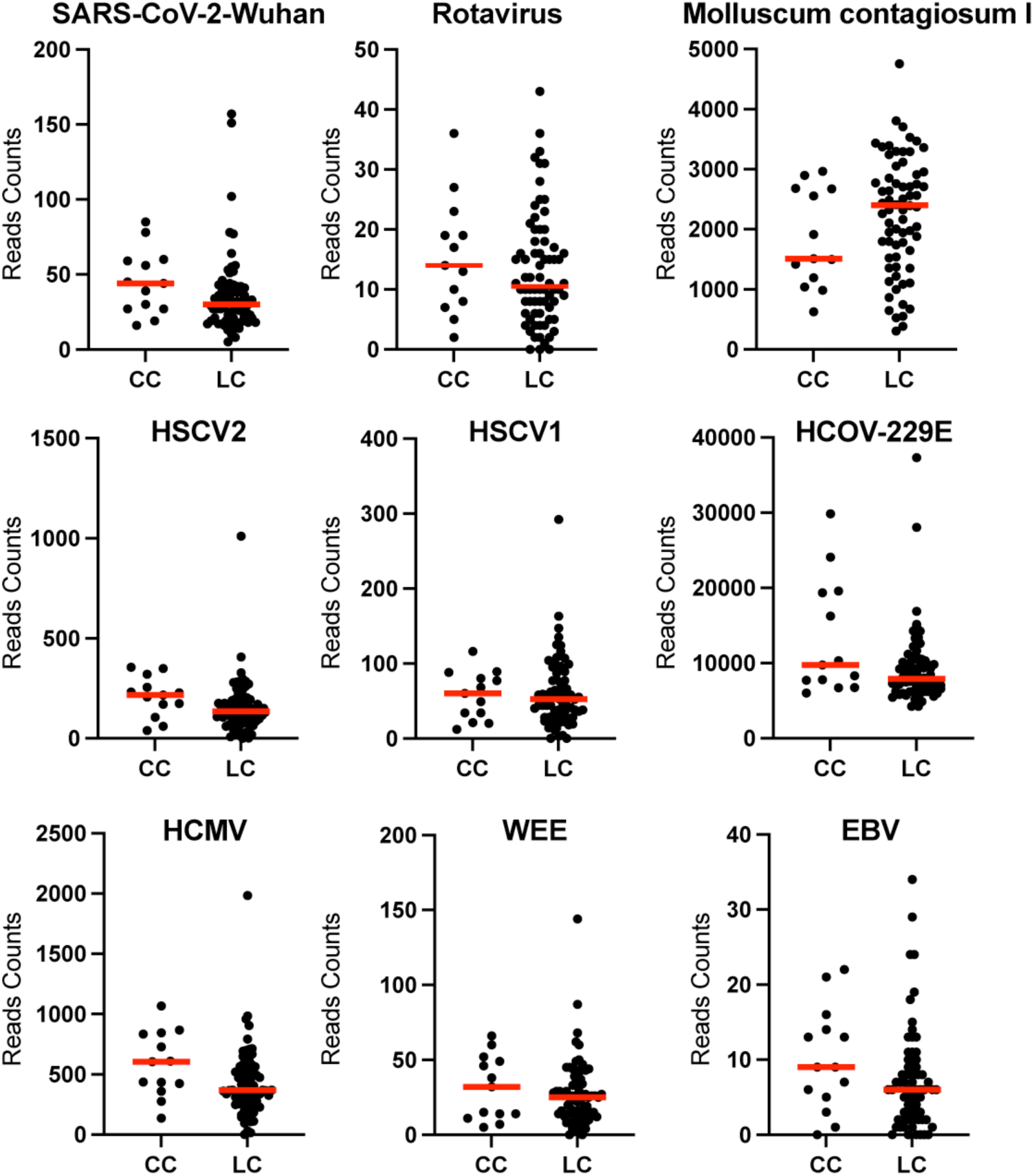
Bulk RNA seq viral reads of six different viruses at 1-12 months after COVID-19 infection in the CC and the LC groups. Enrichment of viral reads in CC and LC groups. Groups comparison was performed using the two-sided Mann-Whitney tests, and P values are shown. ns refers to P>0.05. Medians (red bars) are shown.

**Supplementary Fig. 3.**
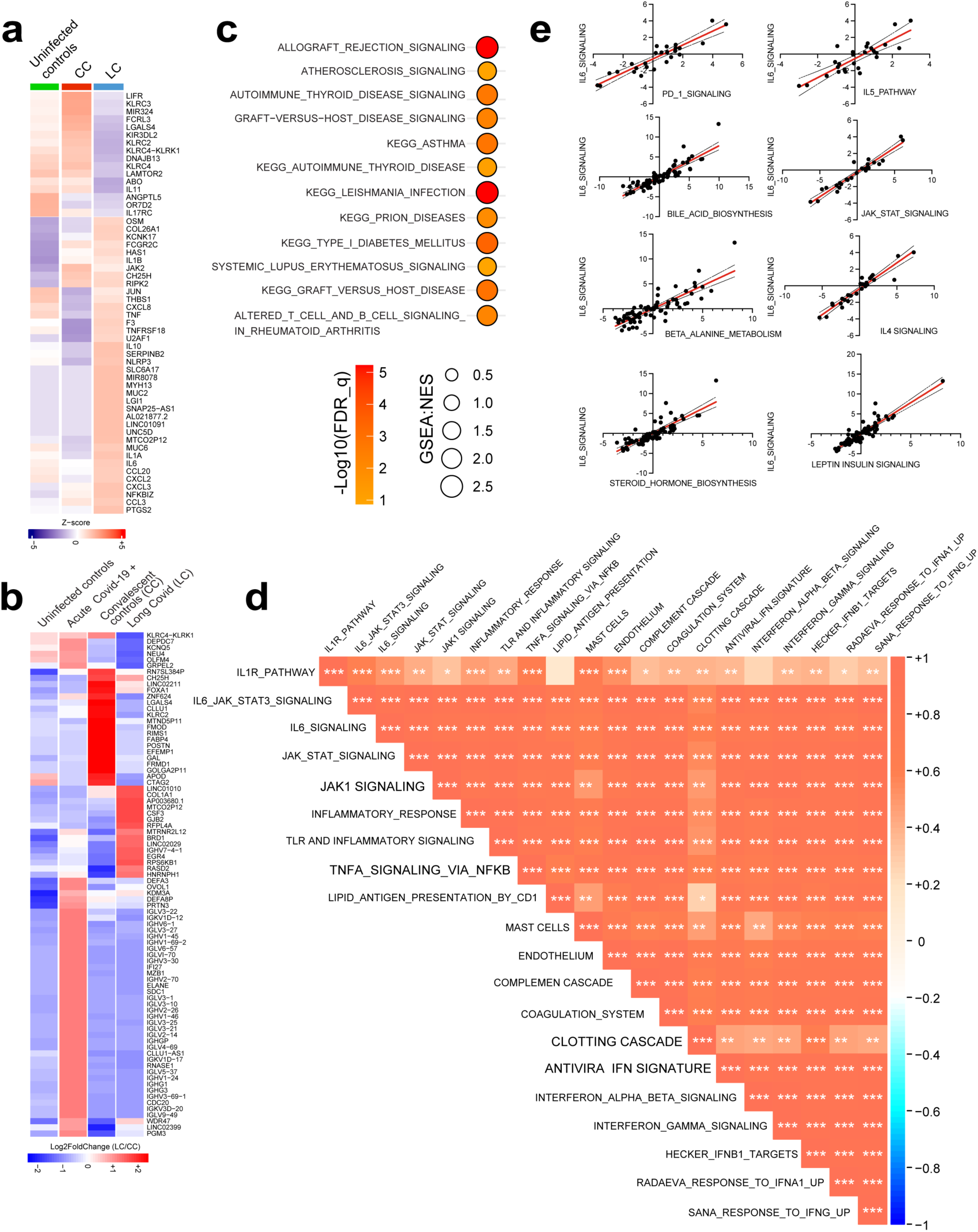
Pathways and genes upregulated (red) or downregulated (blue) in LC compared with CC at 3-6 months to covid19 infection in PBMC. a, Top proinflammatory genes decreased or increased in LC compared with CC and uninfected controls. Heatmap shows the normalized mRNA expression ranging from blue (down regulated) to red (upregulated) in LC group. **b**, log2 fold change expression of the top genes in PBMC (adjusted p value<0.05) in LC vs. CC, LC vs. uninfected and CC vs. uninfected controls. **c,** Circle plots of pathways normalized enrichment scores measured by GSEA of increased pathways in LC compared with CC at 3-6 months to Covid-19 infection. **d**, Correlogram of the pairwise Spearman’s correlation coefficients among multiple signaling pathways involved in inflammatory responses in LC. Each cell in the correlogram shows the correlation between pathways, with the color intensity and the symbol size indicating the strength and direction of the correlation. Positive correlations are depicted towards the red spectrum, while negative correlations are shown in blue, ranging from –1 (perfect negative correlation) to +1 (perfect positive correlation). ***: p-value < 0.001; **: p-value < 0.01; *: p-value < 0.05. **e**, Regression analyses of ssGSEA pathway score of proinflammatory pathways with IL6 signaling. Each point represents an individual sampling time points at 3-6 months after COVID-19 infection. P-values were calculated using two-tailed t-test, indicating the statistical significance of the observed associations.

**Supplementary Fig. 4.**
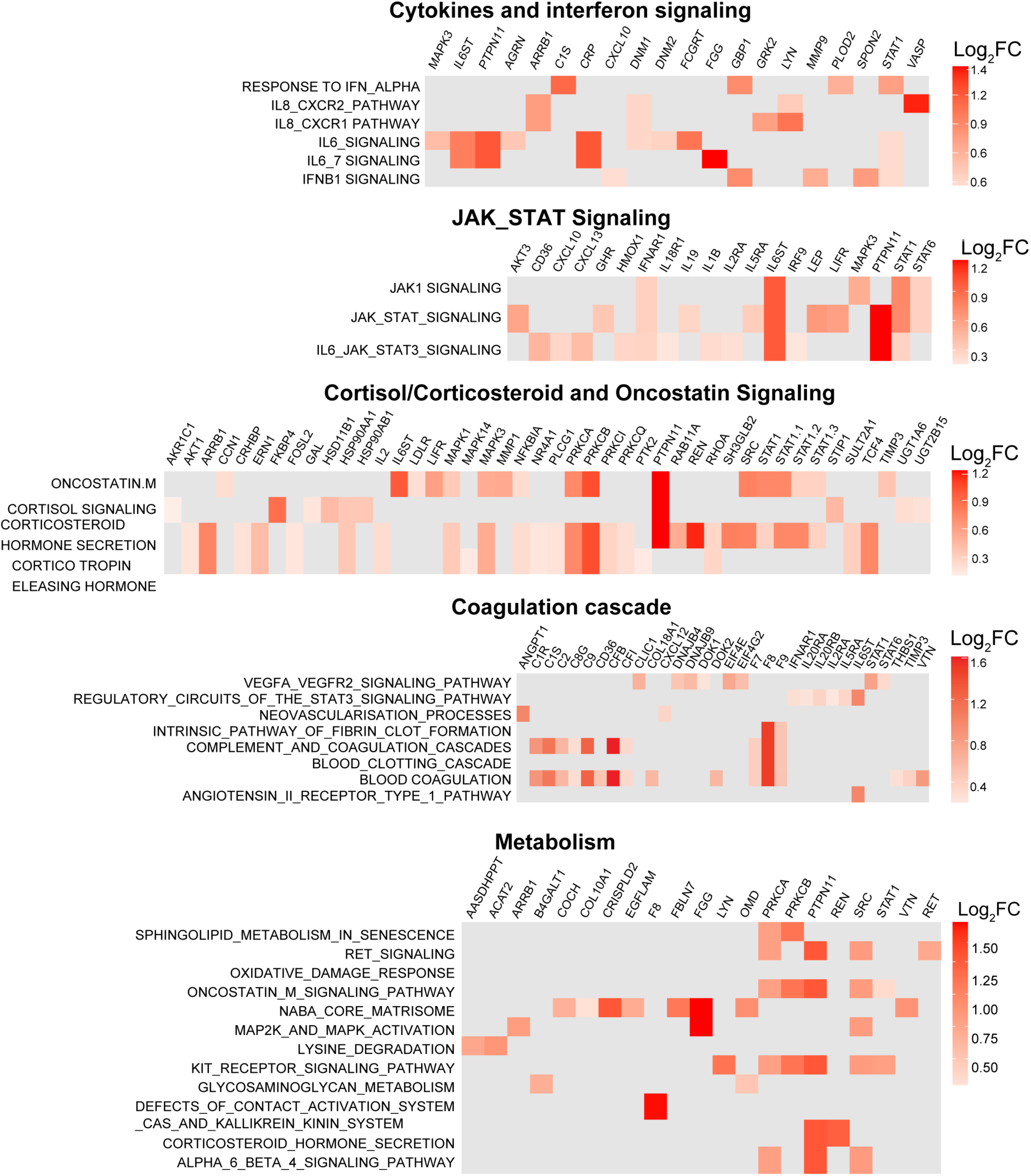
Top proinflammatory proteins increased in LC vs CC at 3_6 months after infection. Increased plasma levels of markers of cytokines and interferon signaling, JAK_STAT signaling, cortisol, oncostatin, coagulation cascade, and metabolism in LC compared with CC patients. Log2 fold change expression in LC vs. CC is shown.

**Supplementary Fig. 5.**
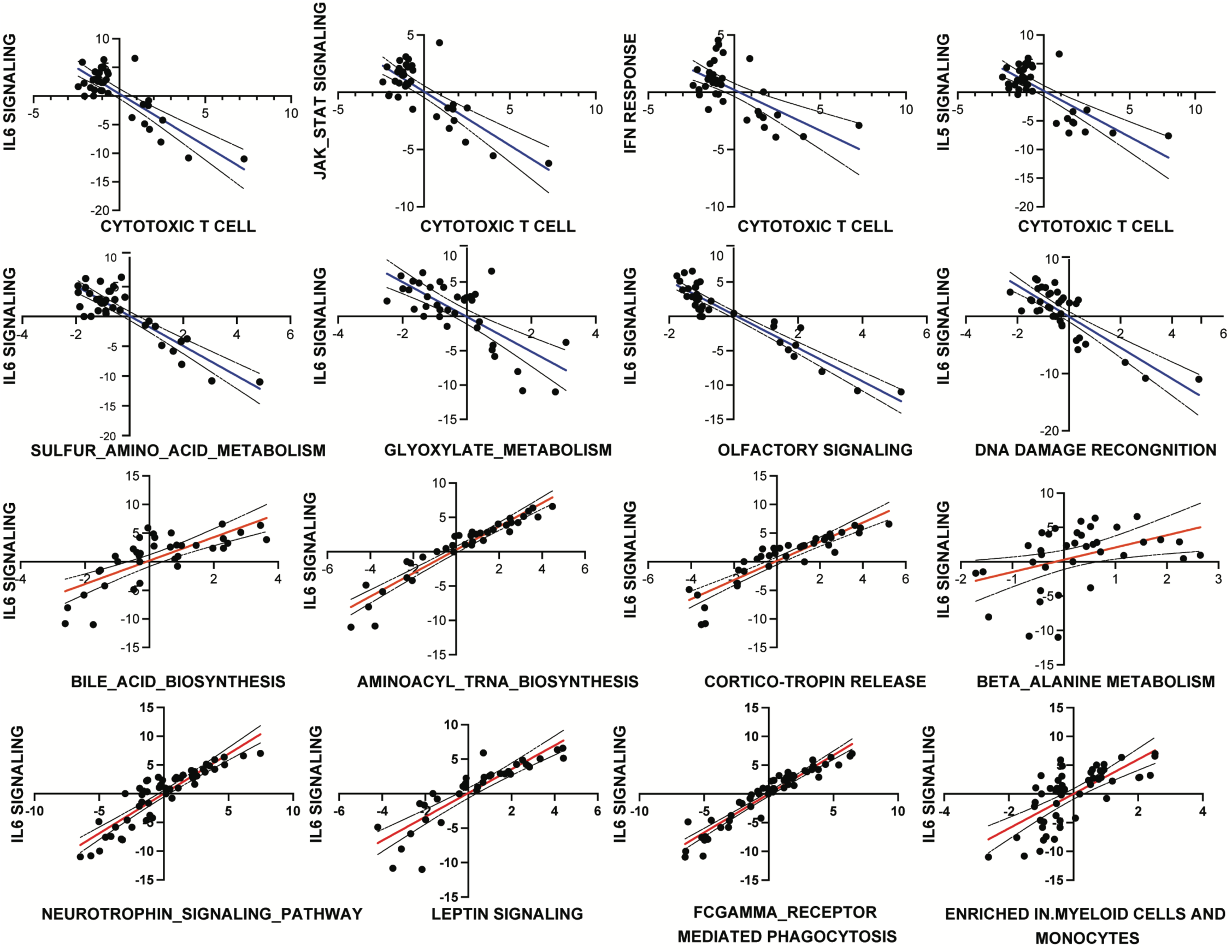
IL6 signaling correlate positively with proinflammatory pathways and immune dysregulation signatures and negatively correlated with cytotoxic T cell signatures in LC. Regression analyses of ssGSEA pathway score of plasma immune signature correlated with plasma IL6 signaling. Each point represents an individual sampling time points at 3-6 months after COVID-19 infection along with corresponding 95% confidence intervals (CI) which illustrate the precision of these estimations. P-values were calculated using two-tailed t-test, indicating the statistical significance of the observed associations (p<0.001).

**Supplementary Fig. 6.**
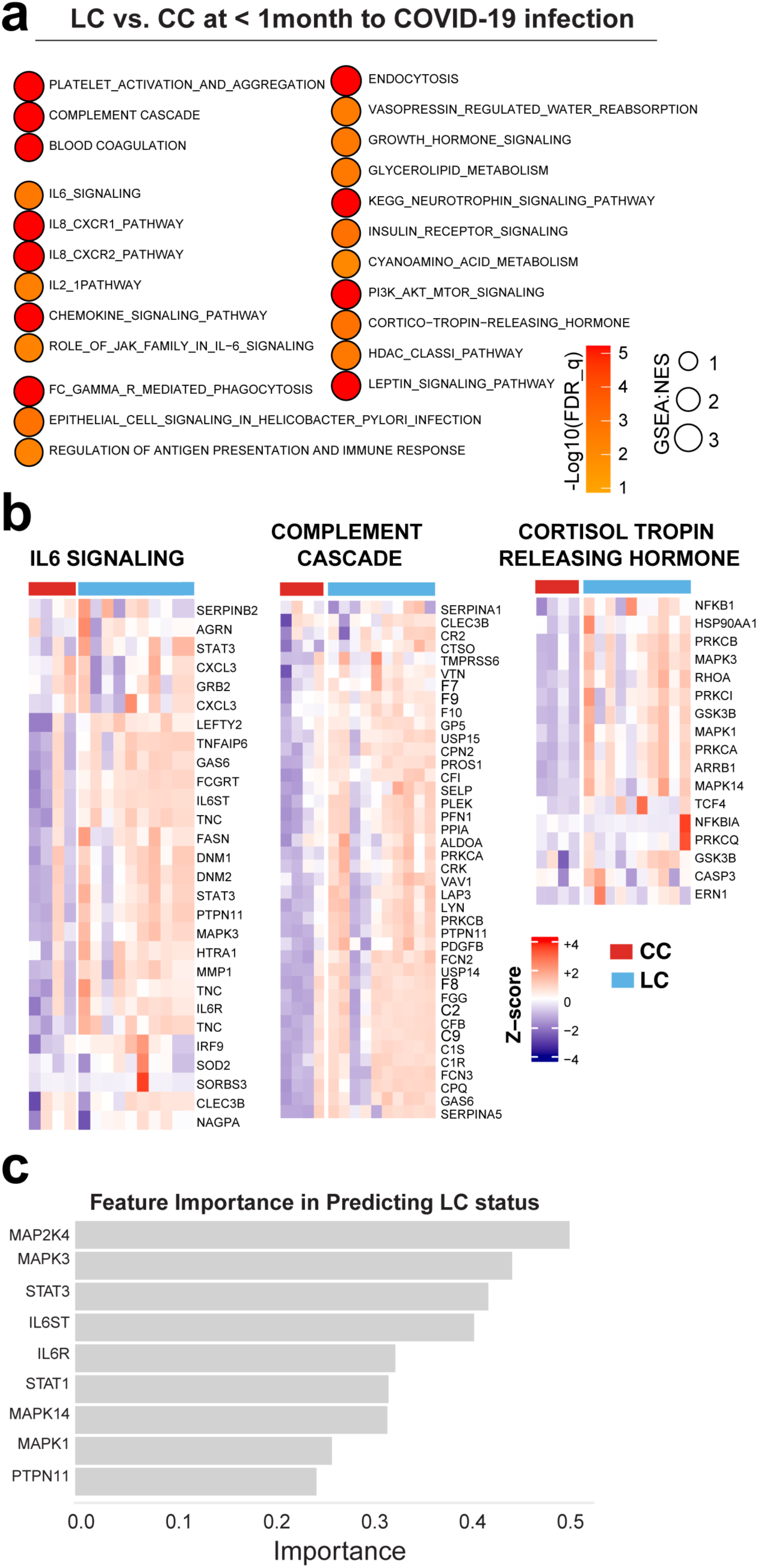
Pathways up or downregulated in LC during the acute phase (<1 month after COVID-19 infection) in plasma. a, Circle plots of pathways normalized enrichment scores measured by gene set enrichment analysis (GSEA, nominal p<0.05). Increased pathways shown in red and decreased pathways shown in blue gradients. Circle size: pathways normalized enrichment score. **b,** Plasma proteomics of IL6 signaling, complement activation and cortisol pathways in LC compared with CC during the acute infection phase (<1mo to COVID-19 infection). Heatmaps of the log 2 (RFU) plasma levels of each protein in LC and CC groups collected within a month after infection is shown. **c**, the top plasma markers during the acute phase predicting the LC status months after infection. Features importance was evaluate using the random forest R package “importance” function and the were sorted using MeanDecreaseGini.

**Supplementary Fig. 7.**
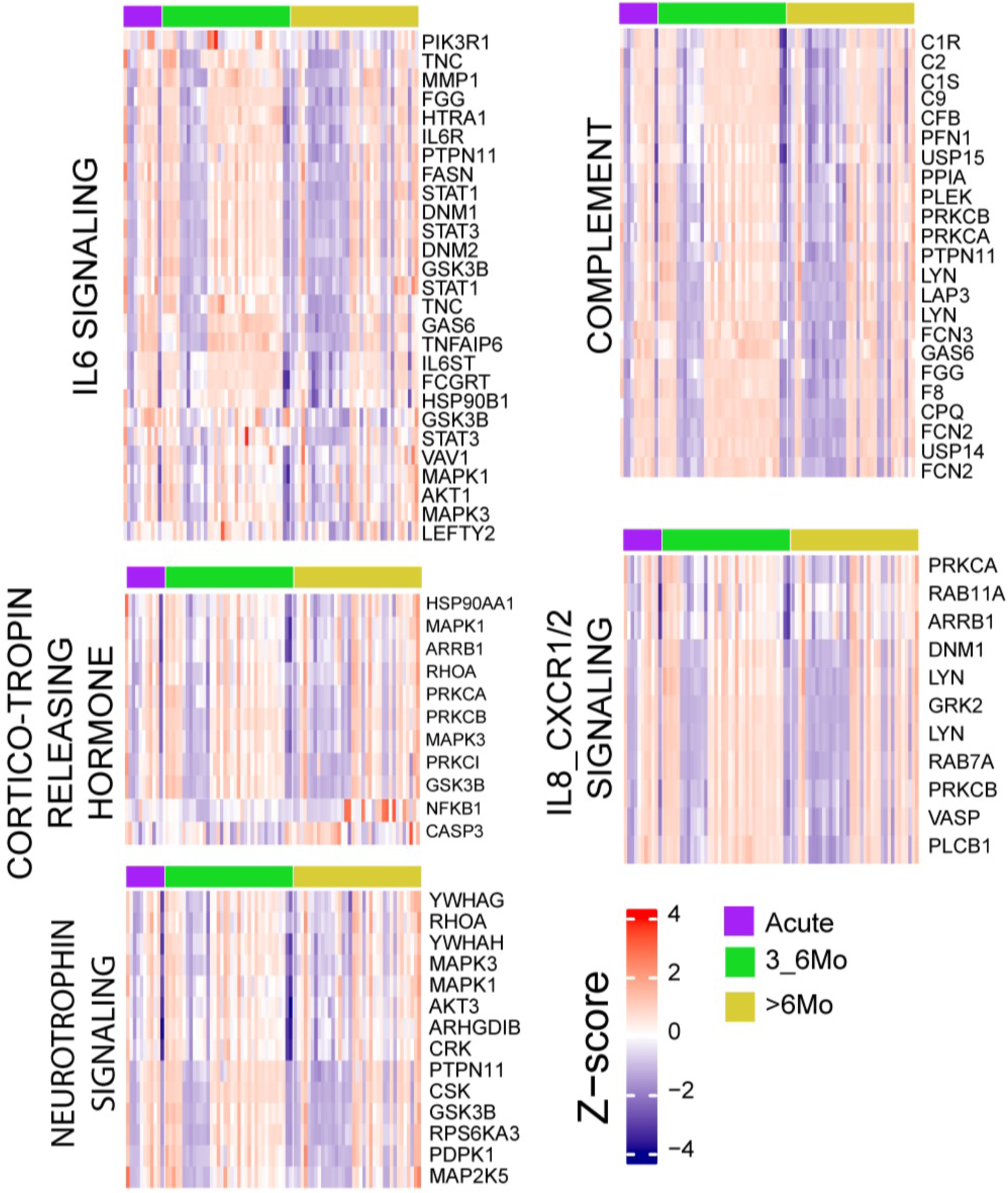
Persistent activation of proinflammatory pathways for more than 6 months after COVID-19 infection in the Long COVID patients. Persistent activation of proinflammatory pathways for more than 6 months after COVID-19 infection in the Long COVID patients. Heatmaps show the normalized plasma levels of the leading genes within each pathway. rows represent genes and columns individual sampling time points ranging from acute infection (< 1 month); 3-6 months and > 6 months after infection in the LC group.

